# Non-parametric analysis of thermal proteome profiles reveals novel drug-binding proteins

**DOI:** 10.1101/373845

**Authors:** Dorothee Childs, Karsten Bach, Holger Franken, Simon Anders, Nils Kurzawa, Marcus Bantscheff, Mikhail Savitski, Wolfgang Huber

## Abstract

Detecting the targets of drugs and other molecules in intact cellular contexts is a major objective in drug discovery and in biology more broadly. Thermal proteome profiling (TPP) pursues this aim at proteome-wide scale by inferring target engagement from its effects on temperature-dependent protein denaturation. However, a key challenge of TPP is the statistical analysis of the measured melting curves with controlled false discovery rates at high proteome coverage and detection power. We present non-parametric analysis of response curves (NPARC), a statistical method for TPP based on functional data analysis and nonlinear regression. We evaluate NPARC on five independent TPP datasets and observe that it is able to detect subtle changes in any region of the melting curves, reliably detects the known targets, and outperforms a melting point-centric, single-parameter fitting approach in terms of specificity and sensitivity. NPARC can be combined with established analysis of variance (ANOVA) statistics and enables flexible, factorial experimental designs and replication levels. To facilitate access to a wide range of users, a freely available software implementation of NPARC is provided.

## Introduction

Determining the cellular interaction partners of drugs and other small molecules remains a key challenge [1, 2, 3, 4]. In drug research, better assays to detect targets (and off-targets) would provide valuable information on drugs’ mechanisms of action, reveal potential reasons for side effects, and elucidate avenues for drug repurposing. More broadly, in cell biology basic research, the dynamical landscape of binding partners of metabolites, messengers or chemical probes contains much uncharted territory. Thermal proteome profiling (TPP) addresses these needs by screening for protein targets of drugs or small molecules in living cells on a proteome-wide scale [5, 6]. TPP combines multiplexed quantitative mass spectrometry with the cellular thermal shift assay (CETSA) [7], which identifies binding events from shifts in protein thermostability (see Figure S1 for a detailed explanation). A typical TPP experiment generates temperature dependent abundance measurements for a large part of the cellular proteome. Drug binding proteins can then be inferred by comparing the melting curves of proteins between samples treated with drug and vehicle (negative control without drug).

Applications of TPP successfully identified previously unknown protein-ligand interactions [5], protein complexes [8] and downstream effects of drugs in signaling networks [6, 9, 10, 11] in human cells. Recently it has also been extended to study drug resistance in bacteria [12] and targets of antimalarial drugs in plasmodium [13]. There is urgent interest in further advancing its component technologies, including experimental and computational aspects, in order to maximize its biological discovery potential [14, 15, 16, 17, 18, 19].

The central computational task in TPP data analysis is the comparison of the temperature dependent abundance measurements—which can be visualized as melting curves—for each protein with and without (or with various concentrations of) drug. The aim is to detect changes in thermostability from statistically significant changes in the melting curves.

A naïve approach is to summarize each curve into a single parameter, such as the melting point (*T*_m_), which is defined as the temperature of half-maximum relative abundance (horizontal line in Figure 1A). Its value is estimated by fitting a parametric model separately for the control and treatment conditions and comparing the estimates. Statistical significance is assessed using replicates and hypothesis testing, such as a *t*- or *z*-test. While the approach has delivered valid and important results [5, 6, 20, 21], we will see in the following that it tends to lead to needlessly high rates of false negatives. There are three main reasons for that: first, drug-induced effects on thermostability do not always imply significant shifts in *T*_m_ (Figure 1B-C). Second, the true *T*_m_ of a protein can lie outside of the measured temperature range, which impairs its estimation (Figure 1D). Both scenarios can result in important targets being missed in the analysis (Figure 1E). The third reason is a more subtle statistical one: hypothesis tests using only the point estimates of *T*_m_ do not take into account goodness-of-fit of the parametric model or the confidence range of the estimates. Thus, important information is ignored, which statistically leads to loss of power.

**Figure 1.**
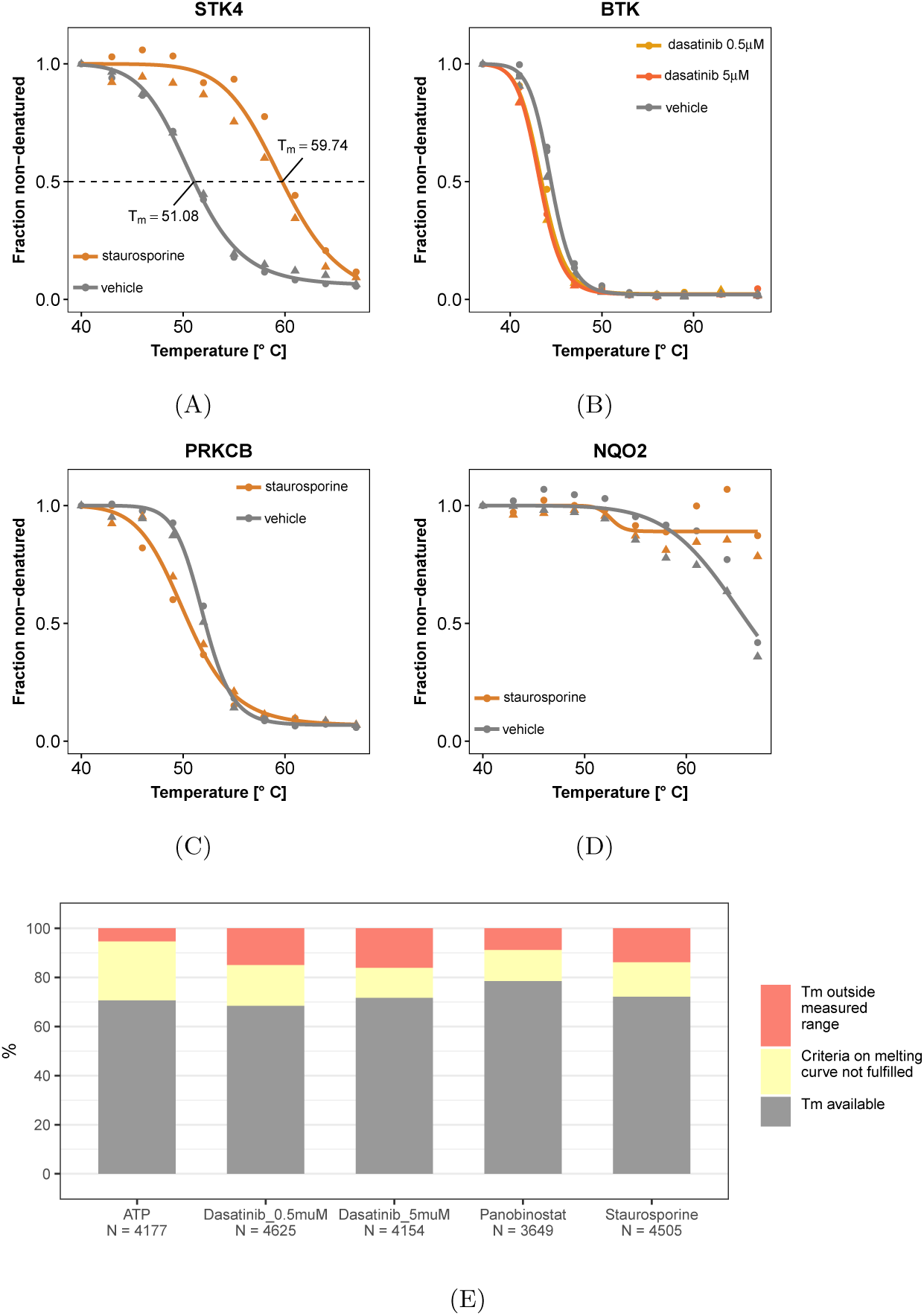
TPP data analysis challenges. **(A)**-**(D)** Examples for protein melting curves with and without drug (see color keys). In each case, ten temperatures were assayed, and two experimental replicates were made per condition, indicated by circle and triangle symbols. Fits of the sigmoid model (Eqn. (3)) to both replicates jointly are shown by smooth lines. **(A)** For serine/threonine protein kinase 4 (STK4), the binding of staurosporine is reflected by a marked shift between the curves. The fitted values for the melting points (*T*_m_) are shown. **(B)** For Bruton’s tyrosine kinase (BTK), there is a small but reproducible shift between the curves. **(C)** Protein kinase C beta (PRKCB) is destabilized by staurosporine; the effect occurs mainly at lower temperatures. **(D)** NAD(P)H quinone dehydrogenase 2 (NQO2) is strongly stabilized by staurosporine. While in each case, the effects of drug binding are clearly reproducible between replicates, the *T*_m_-based approach of [6] only detects (A) and misses (B)-(D). In the case of (B) and (C), the fitted *T*_m_ are too similar, so that the statistical test does not assess the difference as significant. In the case of (D), no reasonable estimate for *T*_m_ in the staurosporine treated condition can be obtained, as it would lie outside the measured temperature range, and the protein is discarded from the analysis. In contrast, NPARC, the method proposed in this article, detects all four cases. **(E)** The fraction of proteins in each of the data sets of Table 1 that is missed by the *T*_m_-based approach due to failure to estimate *T*_m_ or to meet the goodness-of-fit criterion (Table 3, Figure S2).

Here, we propose an alternative approach that compares whole curves instead of summary parameters and does not rely on *T*_m_ estimation. The method, non-parametric analysis of response curves (NPARC), is based on a branch of statistical data analysis that works on continuous functions rather than individual numbers, termed *functional data analysis* [22]. It considers the measured melting curves as samples from an underlying stochastic process with a smooth mean function—which can be modelled parametrically or non-parametrically [23]—and constructs its hypothesis tests directly on these samples. NPARC’s F-statistic uses a more flexible model that makes fewer assumptions on the data than *T*_m_-estimation, is computationally more stable, and it directly uses the information from replicates. As a consequence, reliable estimates of the null distribution of this statistic can be obtained, it shows higher sensitivity for small but reproducible effects, and failures due to model misspecification or outliers are reduced. This increases proteome coverage, which can make the difference between missing or detecting an important drug target.

**Table 1.** Datasets and sample sizes.

| Dataset | Treatment | Concentration | Buffer | Cell line | Intact cells or lysate | Proteins | Reference |
| --- | --- | --- | --- | --- | --- | --- | --- |
| ATP data | MgATP | 2 μM | PBS | K562 | Lysate | 4177 | [9] |
| Dasatinib 0.5 μM data | Dasatinib | 0.5 μM | PBS | K562 | Intact Cells | 4625 | [5] |
| Dasatinib 5 μM data | Dasatinib | 5 μM | PBS | K562 | Intact Cells | 4154 | [5] |
| Panobinostat data | Panobinostat | 1 μM | PBS | K562 | Intact Cells | 3649 | [6] |
| Staurosporine data | Staurosporine | 20 μM | PBS | K562 | Lysate | 4505 | [5] |

**Table 2.**
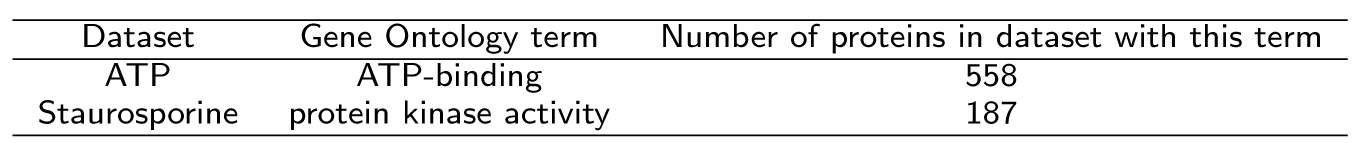
Expected targets per dataset.

**Table 3.** A priori filters applied in the original TPP analysis workflow [6] to select proteins for hypothesis testing.

| Rule number | Rule |
| --- | --- |
| 1 | Both fitted curves for the vehicle and compound treated condition have a coefficient of determination $R^2 > 0.8$, where $R^2 := 1 - \frac{\sum_t (x_t - \mu_c(t))^2}{\sum_t (x_t - \bar{x})^2}$, with $\mu_c(t)$ being the model prediction for condition $c$ at temperature $t$, $\bar{x}$ being the mean of all measurements for the protein within a particular condition and replicate, and the summation extending over all temperatures. |
| 2 | The two curves fitted to the two replicates of the vehicle conditions have a plateau $f_\infty < 0.3$. |
| 3 | In each biological replicate, the steepest slope of the melting curve in the vehicle and treatment condition needs to be $< -0.06$ $^{\circ}\text{C}^{-1}$. |

We demonstrate NPARC on the five published datasets introduced in Table 1. We also compare its results to those of the *T*_m_-based method used by [6]. Three of the experiments used the cancer drugs panobinostat or dasatinib in different concentrations, one investigated the effects of the high-affinity, ATP-competitive pan-kinase inhibitor staurosporine, one the cellular metabolite ATP. While the cancer drugs interact with limited sets of proteins, the two other compounds are promiscuous binders and affect the thermostability of a large fraction of the cellular proteome.

## Experimental Procedures

### Datasets and preprocessing

Five TPP datasets (Table 1) were obtained from the supplements of the respective publications. Each dataset contained relative abundance measurements per protein and temperature which had been scaled to the value measured at 37° C (the lowest of the ten temperatures assayed) and subjected to the global normalization procedure described by Savitski et al. [5]. Only proteins quantified with at least one unique peptide in each of two replicates of the vehicle and compound treated conditions were included in the analysis; the resulting proteome coverages are listed in Table 1.

### Curation of lists of expected targets

Lists of expected protein targets for the pan-kinase inhibitor staurosporine and ATP were obtained from Gene Ontology Consortium annotations via the Bioconductor annotation packages *AnnotationDbi* (version 1.36.2), *org.Hs.eg.db* (version 3.4.0) and *GO.db* (version 3.4.0). Terms and numbers of annotated proteins are shown in Table 2.

### Mathematical model

NPARC is based on fitting two competing models to the data, a null model and an alternative model. The null model states that the relative protein abundance at temperature *t* (given in *°*C) is explained by a single smooth function *µ*_0_ (*t*) irrespective of the treatment condition (Figure 2A). The alternative model posits two condition-specific functions: *µ*_*T*_ (*t*) for the treatment condition and *µ*_*V*_ (*t*) for the vehicle condition (Figure 2B). Deviations between observed data and fitted model are referred to as residuals, and the sum of squared residuals (RSS) serves as an indicator of each model’s goodness-of-fit. We then compute

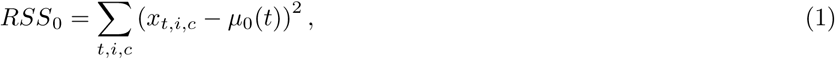

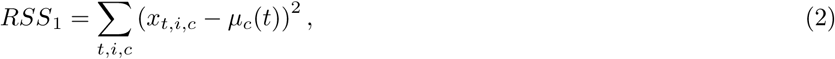

where *x*_*t,i,c*_ is the measured value at temperature *t* for experimental replicate *i* and condition *c ∈* {*V, T*}, and the summations extend over all temperatures, replicates, and conditions.

**Figure 2.**
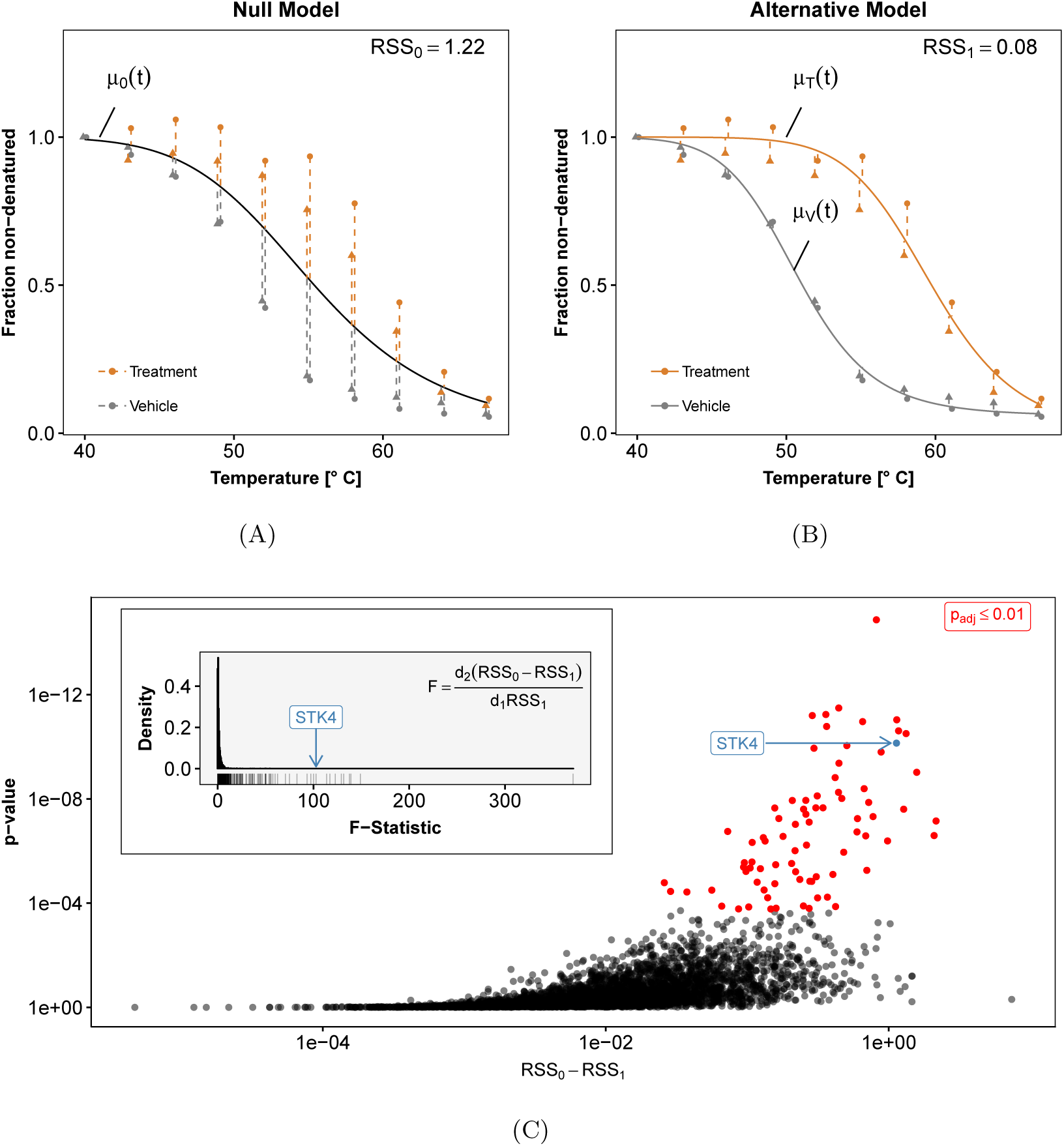
Principles of NPARC, illustrated for protein STK4 under staurosporine treatment. **(A)** Fit of the null model, i.e., no treatment effect (black line). The goodness-of-fit is quantified by *RSS*_0_, the sum of squared residuals (dashed lines). As in Figure 1, the triangle and circle symbols indicate the experimental replicates. **(B)** Fit of the alternative model, with separate curves for the treated (orange) and the vehicle condition (grey). Due to the higher flexibility of the model, the sum of squared residuals *RSS*_1_ is always less than or equal to *RSS*_0_. **(C)** The question whether the improvement in the goodness-of-fit, i.e., the difference *RSS*_0_ – *RSS*_1_, is strong enough to reject the null hypothesis can be addressed with the variant of the *F*-test described in the main text. Each point in the plot corresponds to a different protein. The highlighted example STK4 has a large *F*-statistic and a small p-value.

#### Choice of the mean function

The mean functions *µ*_0_ (*t*), *µ*_*T*_ (*t*) and *µ*_*V*_ (*t*) are each chosen from the same space of smooth functions *f*: ℝ_+_ → [0, 1] spanned by the three parameters *a, b ∈* ℝ, *f*_*∞*_ *∈* [0, 1] and the prescription

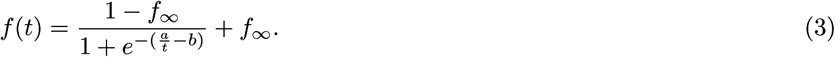

The shape of these functions is sigmoid, and the functional form (3) can be motivated by simplifying protein thermodynamics considerations [5]. The mean functions and the RSS values are computed separately from the data for each protein. In order not to overburden the notation, we omit the protein indices.

### Hypothesis test statistic

To discriminate between null and alternative models, we compute the *F*-statistic

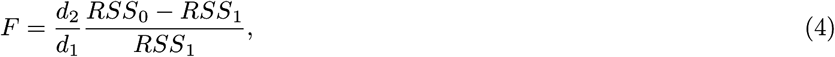

with *d*_2_ */*_1_ > 0 defined as below. *F* quantifies the relative reduction in residuals from null to alternative model. While *F* is by definition always positive, it will be small for proteins not affected by the treatment, while a high value of *F* indicates a reproducible change in thermostability.

#### Null distribution

To compute a p-value from a value of the *F*-statistic (4), we need its null distribution, i.e., its statistical distribution if the data generating process is described by a common mean function *µ*_0_ (*t*). If the residuals were independent and identically normal distributed, this distribution would be given by an analytical formula, namely that of the *F* (*d*_1_, *d*_2_)-distribution with parameters *d*_1_, *d*_2_ > 0, and these parameters—sometimes called *degrees of freedom*—would be explicitly given from the number of measurements and number of model parameters that go into the computation of *RSS*_0_ and *RSS*_1_. In practice, this is not the case, since the residuals are heteroscedastic (i.e. have different variances at different temperatures) and correlated. However, the family of *F* (*d*_1_, *d*_2_)-distributions is quite flexible, and we can approximate the distribution of the *F*-statistic (4) on data occurring in practice with an *F* (*d*_1_, *d*_2_)-distribution with different “effective degrees of freedom” *d*_1_, *d*_2_. To this end, we separately approximate the numerator and denominator of *F* as

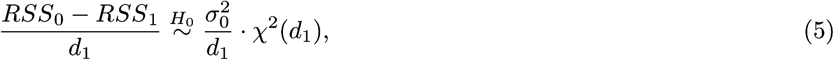

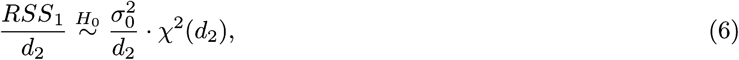

and use the fact that the ratio of two *χ*^2^-distributed random variables *χ*^2^ (*d*_1_), *χ*^2^ (*d*_2_) has an *F* (*d*_1_, *d*_2_)-distribution [24]. The scale parameter 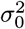 and the effective degrees of freedom *d*_1_ and *d*_2_ are estimated from the the empirical distributions—across proteins—of *RSS*_0_ and *RSS*_1_. Thus, we assume that 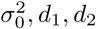, are the same for all proteins. In particular, we estimate 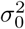 from the moments of *RSS*_1_ − *RSS*_0_ as

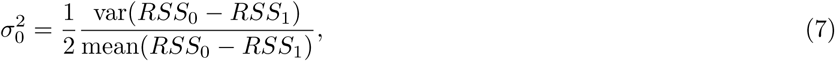

where mean and variance are computed across proteins on the observed values of *RSS*_0_ – *RSS*_1_ (see Supplementary Methods for details). Then, *d*_1_ and *d*_2_ are obtained by numerical optimization of the likelihoods for models (5) and (6) using the `fitdistr` function of the `R` package `MASS` [25].

#### p-values

For each protein, a p-value is computed from its *F*-statistic and the cumulative *F*-distribution with parameters *d*_1_, *d*_2_ as described above. The multiset of p-values across all proteins is corrected for multiple testing with the method of Benjamini and Hochberg [26]. The outcome of such an analysis is exemplarily shown in Figure 2C.

## Results

### Application to panobinostat

We assessed the ability of NPARC to detect drug targets on a dataset on panobinostat (Table 1). Panobinostat is a broad-spectrum histone deacetylase (HDAC) inhibitor known to interact with HDAC1, HDAC2, HDAC6, HDAC8, HDAC10, and tetratricopeptide repeat protein 38 (TTC38) [6].

Out of 3649 proteins reproducibly quantified across both biological replicates in both treatment conditions, NPARC yielded 16 proteins with Benjamini-Hochberg adjusted p-values ≤ 0.01. They contained the expected HDAC targets (Figure 3A-as well as TTC38, the histone proteins H2AFV or H2AFZ (the two variants could not be distinguished by mass spectrometry), and zinc finger FYVE domain-containing protein 28 (ZFYVE28) (Figure 3F-H). These proteins were previously identified as direct or indirect targets of panobinostat [6, 11]. In addition, eight more proteins were detected for which no direct or indirect interactions with panobinostat have been described (Figure S3). They reached statistical significance because they either showed effect sizes comparable to known panobinostat targets, or more subtle but highly reproducible changes in a similar strength to those already described for dasatinib target BTK (Figure 1B). We reanalyzed the more recent 2D-TPP dataset of short-term (15 min) panobinostat-treatment of HepG2 cells [11] for these proteins. All of them were identified and quantified at sufficient peptide coverage, but none of them showed stabilization. We thus conclude that the additionally found proteins are likely not direct binders of panobinostat, but rather indirect effects, like altered protein-protein interactions or post-translational modifications. The longer (5 hours) incubation time of the assay used to generate the panobinostat dataset in Table 1 makes it more sensitive to such effects.

**Figure 3.**
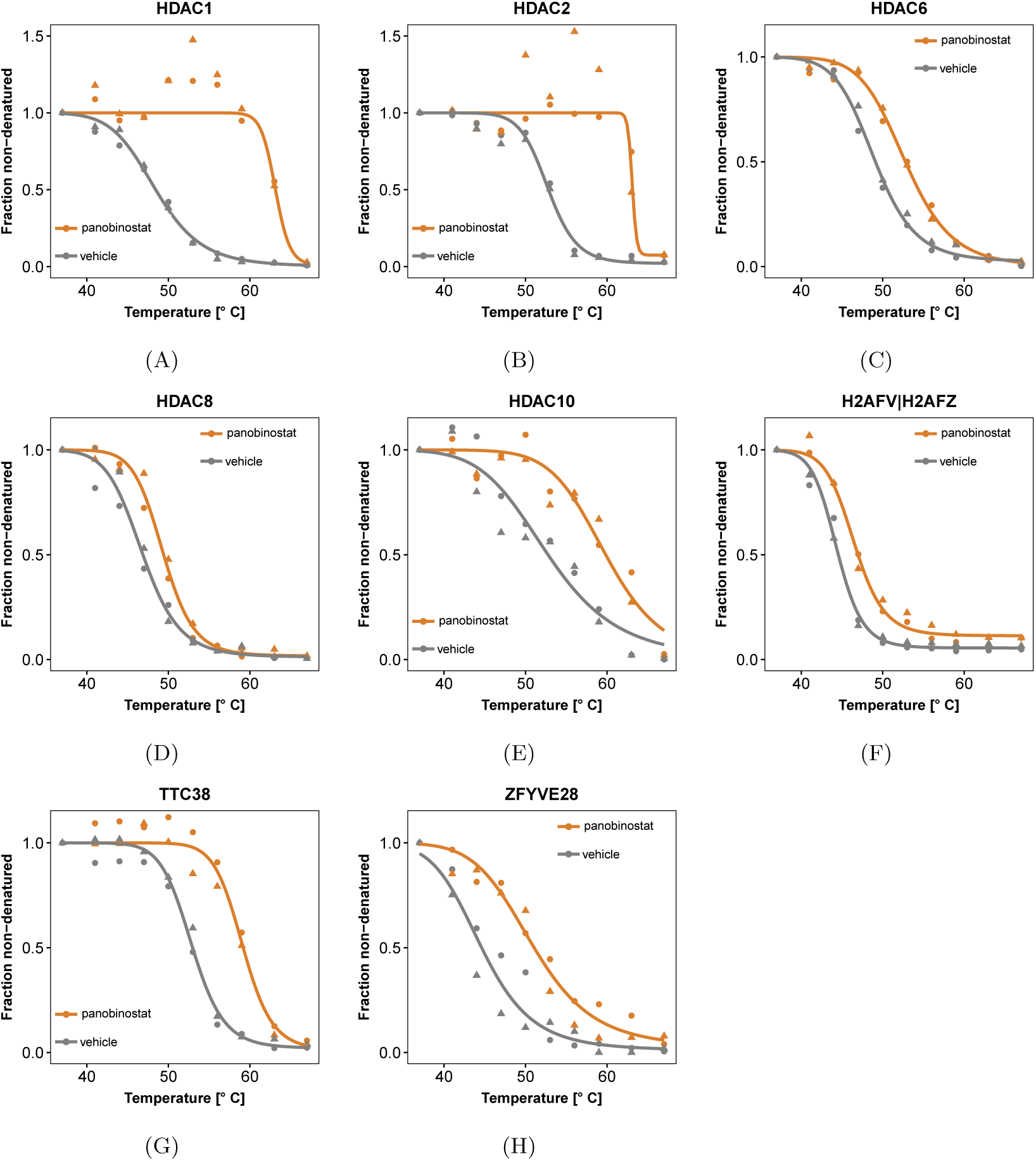
Direct and indirect targets of the HDAC inhibitor panobinostat detected by NPARC. (FDR ≤ 0.01 according to the method of Benjam1ini and Hochberg [26]). **(A)**-**(E)** Data and curve fits for five HDACs that show significant shifts in their thermostability. HDAC1 and HDAC2 are not detected by the *T*_m_-based approach of [6], since the higher variance between the replicates of the panobinostat-treated condition leads to them being eliminated by the filter heuristics of that method (Table 3). In contrast, NPARC naturally takes the variance into account in the computation of the *F*-statistic and does not require such filtering steps. **(F)**-**(H)** Data and curve fits for known non-HDAC targets.

### Beyond two-group comparisons

Since NPARC is based on analysis of variance (ANOVA), it admits experimental designs in which the covariate has multiple levels. An example is the dataset for the BCR-ABL inhibitor dasatinib, which comprises measurements on cells treated at two different concentrations as well as untreated cells. NPARC successfully identified known targets of dasatinib (Figure S4).

### Replicate agreement and model fit diagnostics

Application of the *T*_m_-based approach by [6] to the panobinostat data failed to detect HDAC1 and HDAC2. This was because the data for these proteins had relatively high variance in the drug-treated condition, as is visible in Figure 3A-B. This led to their exclusion according to one of the data quality filter criteria of that method (Table 3), namely the criterion that asks for sufficiently high coefficients of determination (*R*^2^). In contrast, a better and statistically sound trade-off between variability and effect size is an integral part of NPARC, and does not require an *ad hoc* filter criterion.

To further assess the price of the various filter criteria of the *T*_m_-based approach by [6], we tabulated the numbers of proteins affected by them in each of the five datasets. These proteins would, in principle, not be detectable by that method, no matter how strong the effect. Their numbers amounted to 14-27% of the total numbers of proteins for which melting points could be determined in both replicates (Figure 1E and Table 4 and to 21-32% of all proteins irrespective of melting point availability (Figure S2). In contrast, the *F*-test of NPARC could be applied to all proteins irrespective of these or similar criteria, a fact which contributed to the increased protein coverage and sensitivity of NPARC.

**Table 4.** Coverage of proteins applicable for hypothesis testing by the original TPP analysis workflow [6].

| Dataset | $T_m$ outside measured range | $T_m$ available but curves not passing *a priori* filters | $T_m$ available and curves passing *a priori* filters |
| --- | --- | --- | --- |
| ATP data | 220 | 1004 | 2953 |
| Dasatinib 0.5 μM data | 689 | 768 | 3168 |
| Dasatinib 5 μM data | 667 | 507 | 2980 |
| Panobinostat data | 320 | 461 | 2868 |
| Staurosporine data | 621 | 631 | 3253 |

### Effects beyond those on the melting point

Many of the proteins detected by NPARC displayed reproducible changes in curve shape, while their *T*_m_-shifts were small, and not considered significant by the *T*_m_-based approach (Figure 4). An example is the effect of staurosporine on protein kinase C beta (PRKCB), shown in Figure 1C. PRKCB is part of the PKC family, whose members were the first reported staurosporine targets [27, 4] and also exhibit similar characteristics (Figure S5).

**Figure 4.**
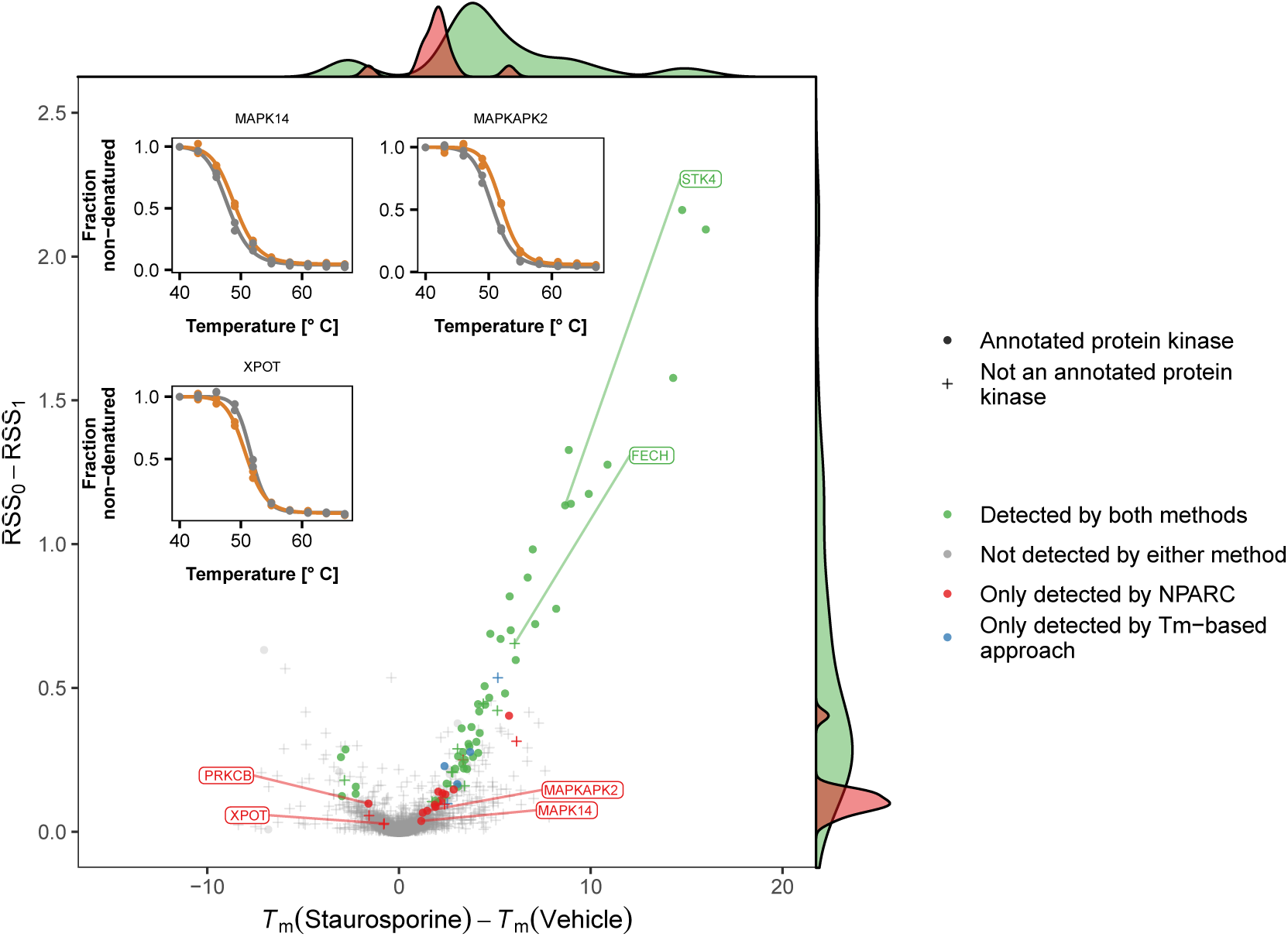
NPARC is sensitive to small but reproducible *T*_m_-shifts. The plot compares the effect size measure used by NPARC, namely *RSS*_0_ – *RSS*_1_ (*y*-axis), to the *T*_m_-difference (*x*-axis) for those proteins in the staurosporine data set for which *T*_m_ estimates could be obtained. Proteins with Benjamini-Hochberg adjusted p-values ≤ 0.01 are marked in red if they were exclusively found by NPARC, and in green if they were also detected by the *T*_m_-based approach of [6]. NPARC detects targets with small *T*_m_-differences if the measurements are reproducible between replicates.

Further examples include the effects of staurosporine on RanGTP binding tRNA export receptor exportin-T (XPOT) and two members of the p38 MAPK signaling pathway: Mitogen-activated protein kinase 14 (MAPK14) and MAP kinase-activated protein kinase 2 (MAPKAPK2) (Figure 4); and the effect of dasatinib on Bruton tyrosine kinase (BTK), an important drug target in B-cell leukemia (Figure 1B).

### Missing melting point estimates

For highly thermostable proteins, the *T*_m_ in one or more of the treatment conditions can be outside of the tested temperature range of a TPP experiment (Figure 1E). One example is NAD(P)H quinone dehydrogenase 2 (NQO2), a cytosolic flavoprotein and a common off-target of kinase inhibitors [28, 29, 30]. In concordance with previous CETSA studies that found NQO2 to be highly stable [31], we observed denaturation only beginning at 67 *°*C (Figure 1D). Staurosporine treatment further stabilized NQO2 to an extent that it showed no sign of melting in the tested temperature range. The *T*_m_-based approach by [6] will discard such proteins, in order to avoid potential problems from extrapolation of the fit beyond the measured temperature range. In contrast, the functional data analysis approach of NPARC is able to detect changes in any part of the melting curves, without reference to a single point such as *T*_m_.

### Sensitivity and specificity

So far, we have described increased sensitivity of NPARC, i.e., its ability to detect more true targets. However, this is only useful if at the same time specificity is maintained, i.e., if false positive detection remains under control. To compare these performance characteristics between NPARC and the *T*_m_-based approach, we computed pseudo receiver operator characteristic (ROC) curves for each of these methods on the staurosporine data and the ATP data, using as pseudo ground truth lists of expected targets from Gene Ontology annotation (Table 2). Here, the term *pseudo* refers to the fact that these target lists, and hence the ROC curves, are only approximations of the truth; however, the relative ranking of two methods in such a pseudo-ROC comparison is likely to be faithful even in the presence of such approximation error [32].

Figure 5 shows the results of NPARC on both datasets, as well as those of the *z*- test of the *T*_m_-based approach by [6] applied to the individual replicates (displayed as a continuous lines parameterized by the *z*-cutoff), and those of the full procedure of [6] (shown by isolated points, due to its single, fixed cutoff). On the staurosporine data, the full procedure of [6] performs close to NPARC. For the individual *z*-tests, as well as overall on the ATP data, NPARC shows superior performance. Given that the decision rule set of [6] (listed in Table 5) and its cutoff parameters were developed and tuned partly on the staurosporine data, these results indicate that NPARC has fewer “fudge parameters” and is likely to be superior in applications to new datasets.

**Table 5.**
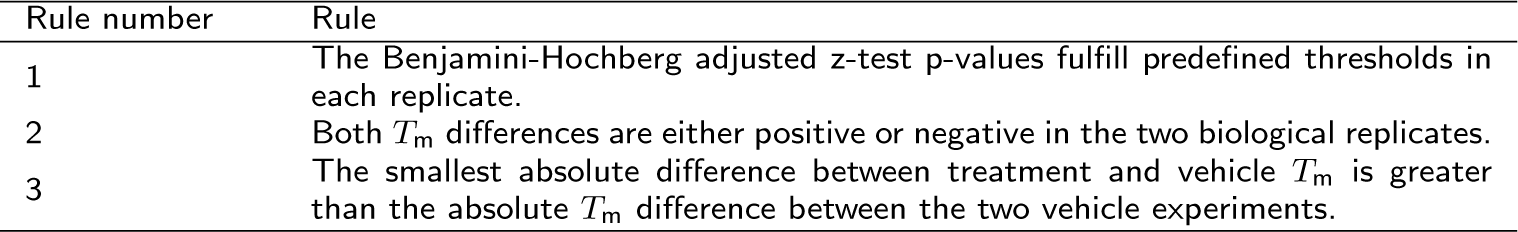
Decision ruleset applied in the original TPP analysis workflow [6] to combine z-test p-values across replicates in an experimental design with two biological replicates.

**Figure 5.**
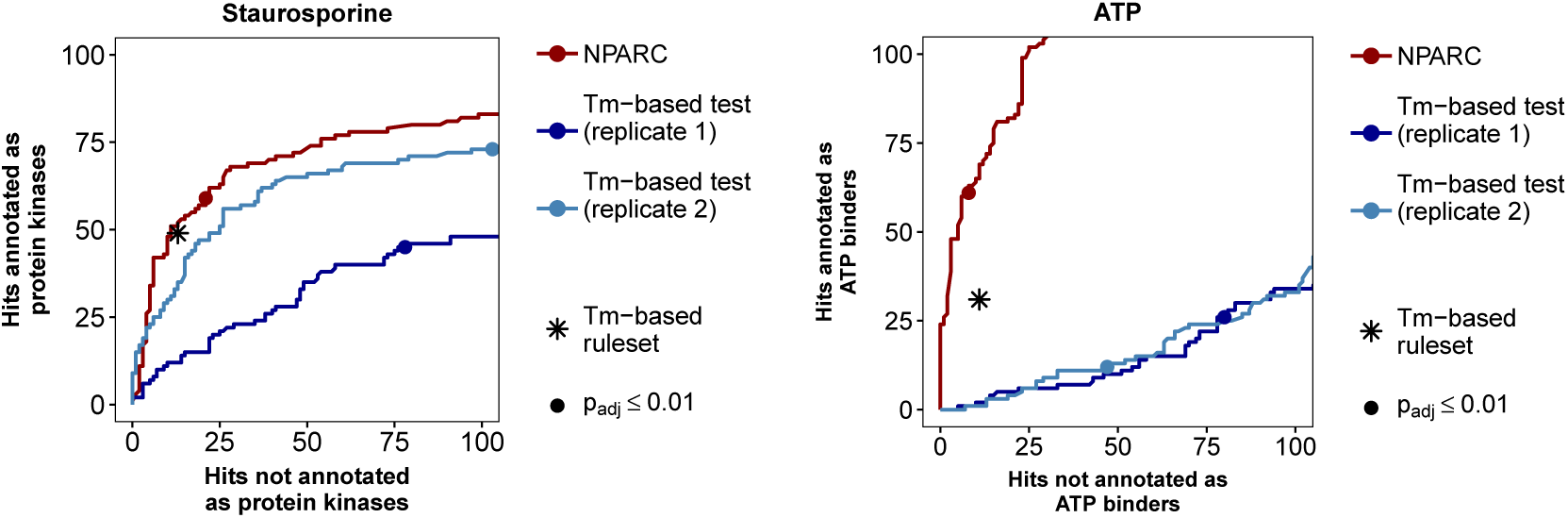
Sensitivity and specificity. Shown are pseudo-ROC curves, with expected hits (as a proxy for true positives) along the *y*-axis and unexpected hits (as a proxy for false positives) along the *x*-axis. The curves are obtained by varying the p-value cutoff of the *F*-test of NPARC (which is computed across replicates), and of the *z*-test of the *T*_m_-based approach [6] (which is computed separately for each replicate). The asterisks indicate the result from the decision rules of [6] on the *z*-test results (Table 5). The dots indicate a threshold of 0.01 on the Benjamini-Hochberg adjusted p-values from NPARC (derived on both replicates in parallel) and on the Benjamini-Hochberg adjusted p-values from the T_m_-based approach (computed individually for each replicate). NPARC is modestly better than the *T*_m_-based approach on the staurosporine data **(A)**, and substantially better on the ATP data **(B)**. The proteins found by NPARC at Benjamini-Hochberg adjusted p-value ≤ 0.01 are also shown in Supplementary Figures S6 and S7.

## Discussion

Thermal proteome profiling offers the possibility to comprehensively characterize ligand-protein interactions on a proteome-wide scale in living cells. However, the method poses the analytical challenge of how to identify statistically significant shifts in thermostability amongst thousands of measurements.

To address this challenge, we introduced a functional data analysis approach to test for treatment effects by comparing competing models by their goodness-of-fit. This enables detection of treatment effects even if a (de-)stabilization of a protein is not captured by a single summary parameter like the *T*_m_. The presented method is based on a sound statistical foundation and does not rely on hard-to-choose cutoff or tuning parameters. We showed that our method compares favourably to previous approaches with respect to sensitivity and specificity for several exemplary datasets, including ones with specific and ones with promiscuous binders.

The approach fits into the framework of analysis of variance (ANOVA) or linear models, and can thus be extended to experimental designs more complex than treatment-control comparisons, such as multiple levels (e.g. drug concentrations) per covariate, multiple covariates and interactions.

The suggested framework is flexible with regard to the mean function used to represent the melting behavior and can be adapted to the particular biological process of interest. To represent nonlinear relationships, approaches include locally linear regression [33], spline regression [34, 35] and nonlinear parametric regression. Here, we chose the latter as it incorporates *a priori* knowledge about the data and thus has favourable estimation efficiency. For example, sigmoid curves have horizontal asymptotes at both sides of the temperature range. In contrast, splines and local regression tend to overfit data near the boundaries of the observation range.

In a cellular environment we occasionally observe non-sigmoid melting curves for subsets of proteins. One possible reason is the presence of protein subpopulations each with distinct melting curves [16]. For example, the formation of protein complexes, the binding to other molecules, or the localization in cellular compartments can lead to deviations from the idealized sigmoid melting curve expected from the same protein in purified form. Our model currently does not account for such systematic and reproducible shape deviations. This could be adapted in future work by adding a low-parametric systematic modification to the sigmoid mean function. We have considered CETSA experimental designs, where the temperature is the major experimental variable and drug concentration is either zero or a chosen value. It appears relatively straightforward to extend NPARC to the isothermal dose response (ITDR) design [6, 7] where temperature is held constant and the drug concentration is varied across a range of values. A further extension of interest would be to 2D-TPP [11] where both factors are changed.

We employ the same “average” null distribution for all proteins, which we obtain by estimating its parameters (*d*_1_, *d*_2_, *σ*_0_) from the distributions of residuals across all proteins. It is conceivable that determining null distributions in a protein-dependent manner, for instance by stratification, could increase the overall power of the method.

The here presented approach is likely to increase the accuracy of profiling protein-ligand interactions in living cells. To facilitate access to a wide range of users we have implemented NPARC in a freely available R workflow [36].

### Additional Files

The following Figures and Tables can be found in the Supplementary Material.

Additional file 1 — Supplementary Methods

Detailed description of the fitting procedures for the scaling parameter, melting points, and mean functions of the model.

Additional file 2 — Table S1

Spreadsheet containing the results of the NPARC approach and of the *T*_m_-based approach for all datasets listed in Table 1.

Additional file 3 — Supplementary Figures S1-S5

PDF containing Supplementary Figures S1-S5.

Additional file 4 — Supplementary Figure S6

All proteins detected by the NPARC approach with Benjamini-Hochberg adjusted *F*-test p-values ≤ 0.01 in the staurosporine data.

Additional file 5 — Supplementary Figure S7

All proteins detected by the NPARC approach with Benjamini-Hochberg adjusted *F*-test p-values ≤ 0.01 in the ATP data.

## Abbreviations

CETSA: Cellular thermal shift assay
FDR: False discovery rate
H_0_: Null hypothesis
H_1_: Alternative hypothesis
NPARC: Non-parametric analysis of response curves
ROC: Receiver operating characteristic
RSS: Residual sum of squares
T_m_: Melting point
TPP: Thermal proteome profiling

## Acknowledgments

We would like to thank Sindhuja Sridharan (EMBL Heidelberg) for help with Figure preparation. We thank EMBL and GlaxoSmithKline for supporting the work. KB is supported by a Cambridge Cancer Centre studentship. SA is funded by the Deutsche Forschungsgemeinschaft, SFB 1036. WH acknowledges funding from the European Commission’s H2020 Programme, Collaborative research project SOUND (Grant Agreement no 633974).

## Software and data availability

The TPP-TR experiments based on staurosporine-, panobinostat-, ATP-, and dasatinib treatments are included in the supplementary materials of references [5, 9, 6]. All results generated from this data are provided in the supplementary material attached to this work. A vignette describing the workflow and relevant code to reproduce the results is available for download [36]. An open-source R/Bioconductor package `NPARC` will soon be released.

## Authors’ contributions

KB, DC and HF contributed equally. KB, DC, HF, SA, MS and WH developed the model. KB and DC implemented the software package and the analytical computations. KB, DC, HF, NK, MS, MB and WH interpreted the results. KB, DC, HF, NK, SA and WH wrote the manuscript. All authors read and approved the final manuscript.

