## Supplementary Methods for "Non-parametric analysis of thermal proteome profiles reveals novel drug-binding proteins"

September 1, 2019

### Estimating the scaling parameter $\sigma_0^2$

The scaled $\chi^2$-distribution in Eqn. (5) of the main text can be alternatively expressed as a $\Gamma$-distribution with shape parameter

$$\alpha = \frac{d_1}{2} \tag{1}$$

and scale parameter

$$\theta = 2\sigma_0^2. \tag{2}$$

The shape and scale parameters are related to the mean $M$ and variance $V$ of the $\Gamma$-distribution as follows:

$$M = \alpha \cdot \theta \tag{3}$$
$$V = \alpha \cdot \theta^2. \tag{4}$$

Substituting (1) and (2) into (3) and (4) yields expressions for $M$ and $V$ in terms of $d_1$ and $\sigma_0$:

$$M = \sigma_0^2 \cdot d_1 \tag{5}$$
$$V = 2 \cdot \sigma_0^4 \cdot d_1. \tag{6}$$

Solving (5) and (6) for $d_1$ yields

$$d_1 = 2 \cdot \frac{M^2}{V}. \tag{7}$$

and for the scaling factor $\sigma_0^2$:

$$\sigma_0^2 = \frac{\theta}{2} = \frac{M}{2\alpha} = \frac{M}{d_1} = \frac{M}{2 \cdot \frac{M^2}{V}} = \frac{1}{2}\frac{V}{M}. \tag{8}$$

These equations coincide with those used by Brown's method [1], which proposes an adaptation of Fisher's method for combining multiple p-values to the scenario of correlated tests by estimating $\chi^2$-distribution parameters from the data in a similar manner. To increase robustness, we estimated $M$ and $V$ by the median and median absolute deviation (also called *D-estimates* by [2]) of the observed values of $RSS_1 - RSS_0$.

### Model fitting

All mean functions were fitted by nonlinear least squares regression using the `nls` function in `R`. For the NPARC analysis, the melting curve model (Eqn. (3) in the main text) was fitted separately per protein to obtain $\mu(t)$, or per protein and treatment condition to obtain $\mu_c(t)$. To reproduce the results of the $T_\mathrm{m}$-based approach (Figure 5), the model fits were repeated per replicate and treatment condition for each protein.

### Summary of the $T_\mathrm{m}$-based approach

The results of the $T_\mathrm{m}$-based approach were obtained with the `R` package `TPP` [3]. This package, and the method it implements, are described in [4], and we only briefly summarize here. For each curve obtained by the replicate- and condition-wise model fits, $T_\mathrm{m}$ was calculated as $T_\mathrm{m} = a/(b - \ln(\frac{1-f_\infty}{0.5-f_\infty} - 1))$ so that it fulfilled $f(T_\mathrm{m}) = 0.5$. Before hypothesis testing on this parameter, *a priori* filters were applied to remove curves with undesirable shape or goodness-of-fit by setting a threshold on the $R^2$, the slope and the plateau parameters (Table 3). Within each replicate, the difference in $T_\mathrm{m}$ between the treatment and control conditions ($\Delta T_\mathrm{m}$) was computed per protein and converted to z-scores. Robust versions of the z-scores were computed by replacing the mean and standard deviation by quantiles of the empirical distributions of $\Delta T_\mathrm{m}$, namely the mean by the 0.5 quantile, and the standard deviation by the 0.8413 quantile for positive values, or the 0.1587 quantile for negative values. These quantities correspond to mean and standard deviation in the case of a normal distribution. In order to reduce the influence of values with high estimation uncertainty when calculating these quantiles, proteins were binned by the slopes of their curves, and z-scores were calculated separately for each bin as described in [5]. For each protein, p-values were calculated by comparing the z-scores to the normal distribution. To reach the final decision for each protein, the p-values were combined heuristically across replicates using the decision ruleset of Table 5.
