## Supplementary Figures S1-S5 for "Non-parametric analysis of thermal proteome profiles reveals novel drug-binding proteins"

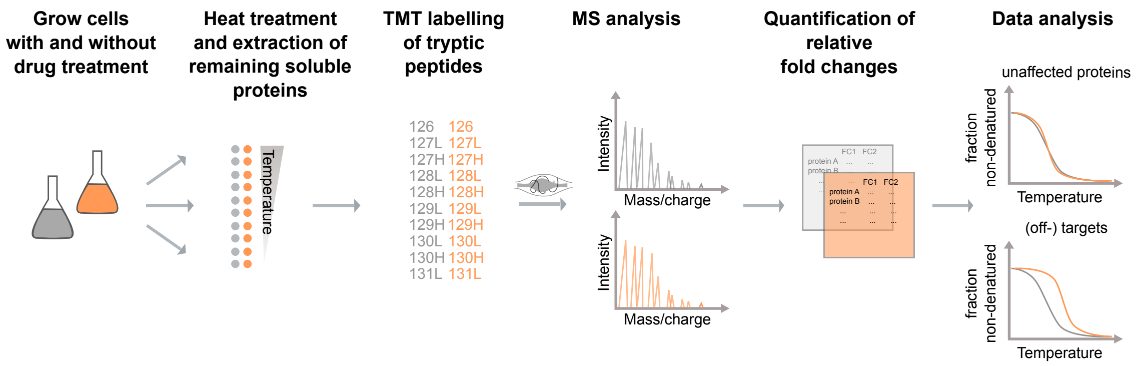


Figure S1. Experimental workflow of TPP experiments. After ligand (orange) or vehicle (grey) treatment, the cells or cell extracts are split up into ten aliquots and heated to different temperatures. This is followed by protein extraction, trypsin digestion and labeling with TMT10 isobaric tags specific to each temperature. Protein abundances are quantified by the reporter ion intensities in the LC-MS/MS. Finally, fold changes are computed that reflect the protein concentration relative to the lowest temperature.

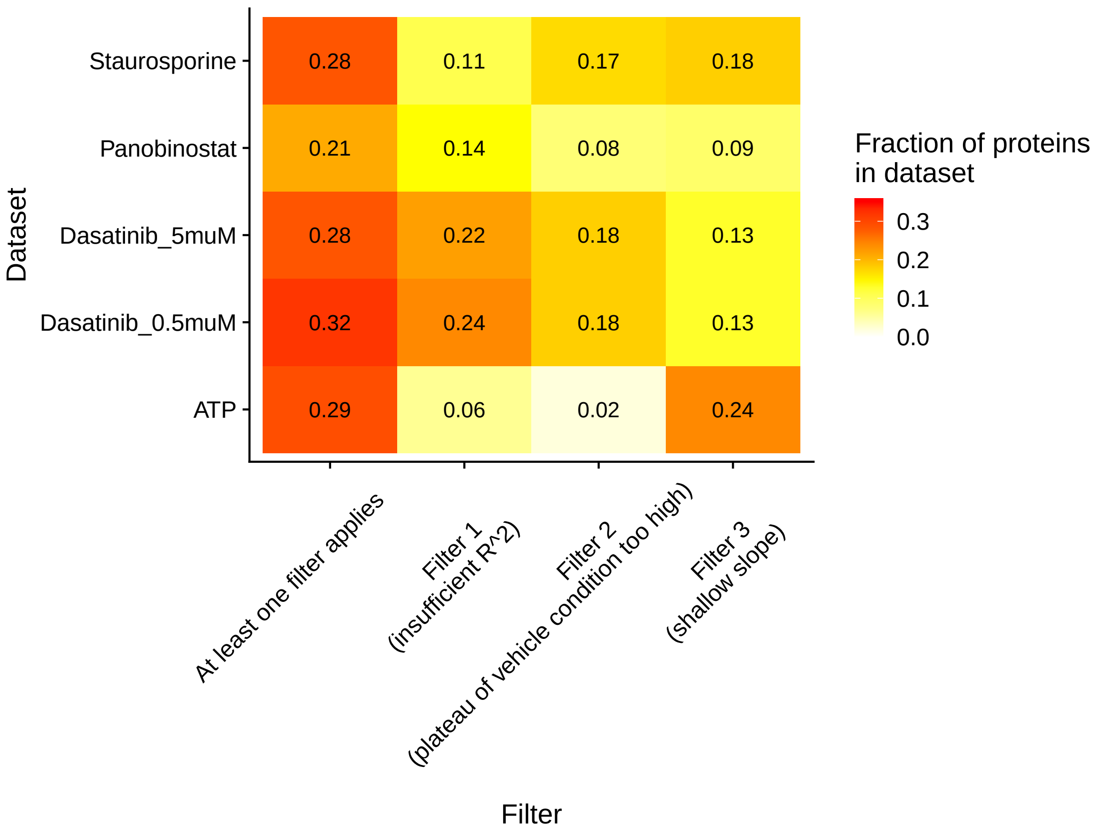


Figure S2. Effects of QC-filters (Table 3) applied to restrict the analysis to 'well-behaved' curves for hypothesis testing in the $T_{\mathrm{m}}$-based approach of [6]. The numbers and colors indicate the fraction of proteins in each dataset that is removed from the analysis by each filter.

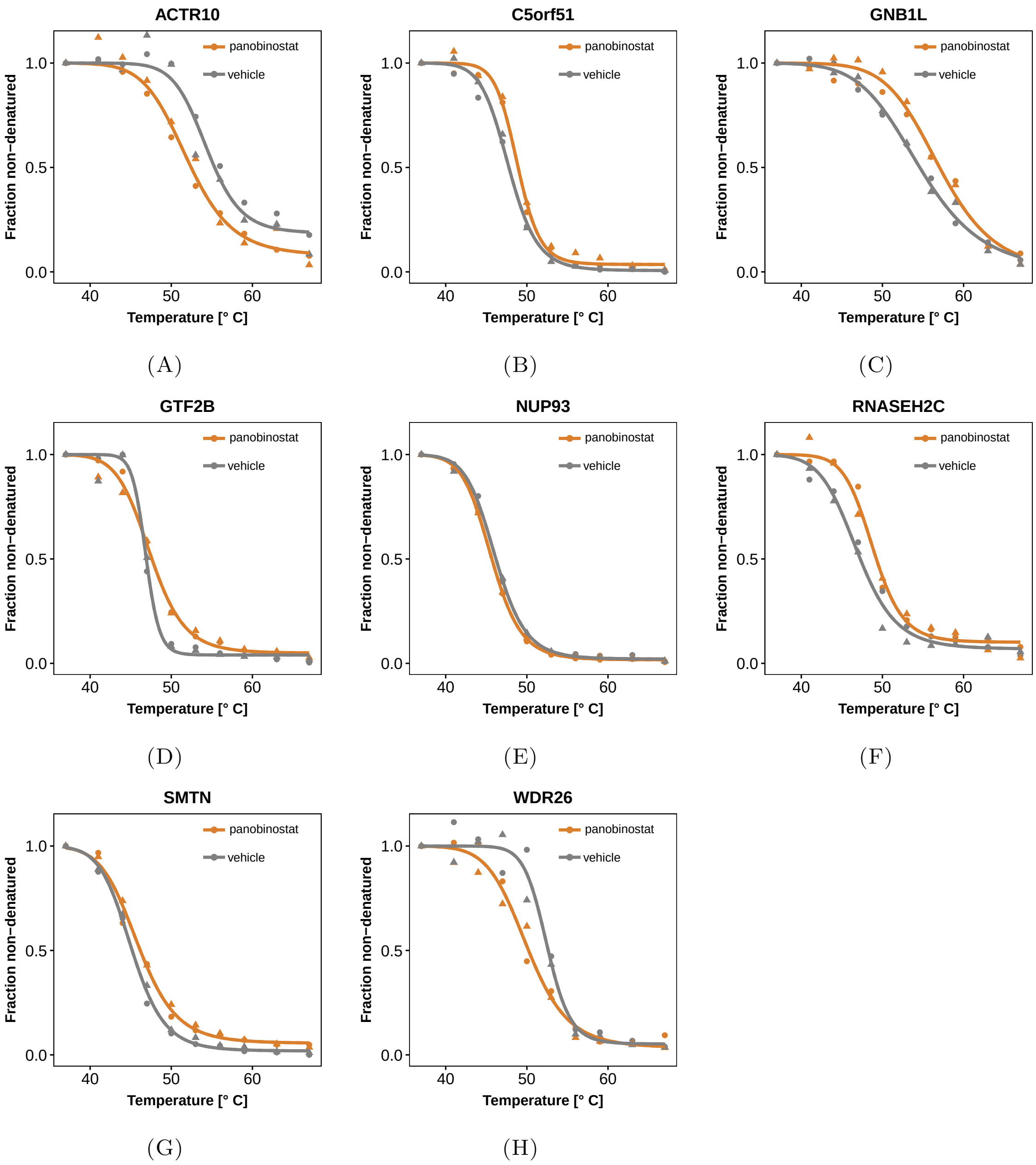


Figure S3. Melting curves of the eight panobinostat hits detected with Benjamini-Hochberg adjusted p-values $\leq 0.01$ in addition to the expected targets in Figure 1.

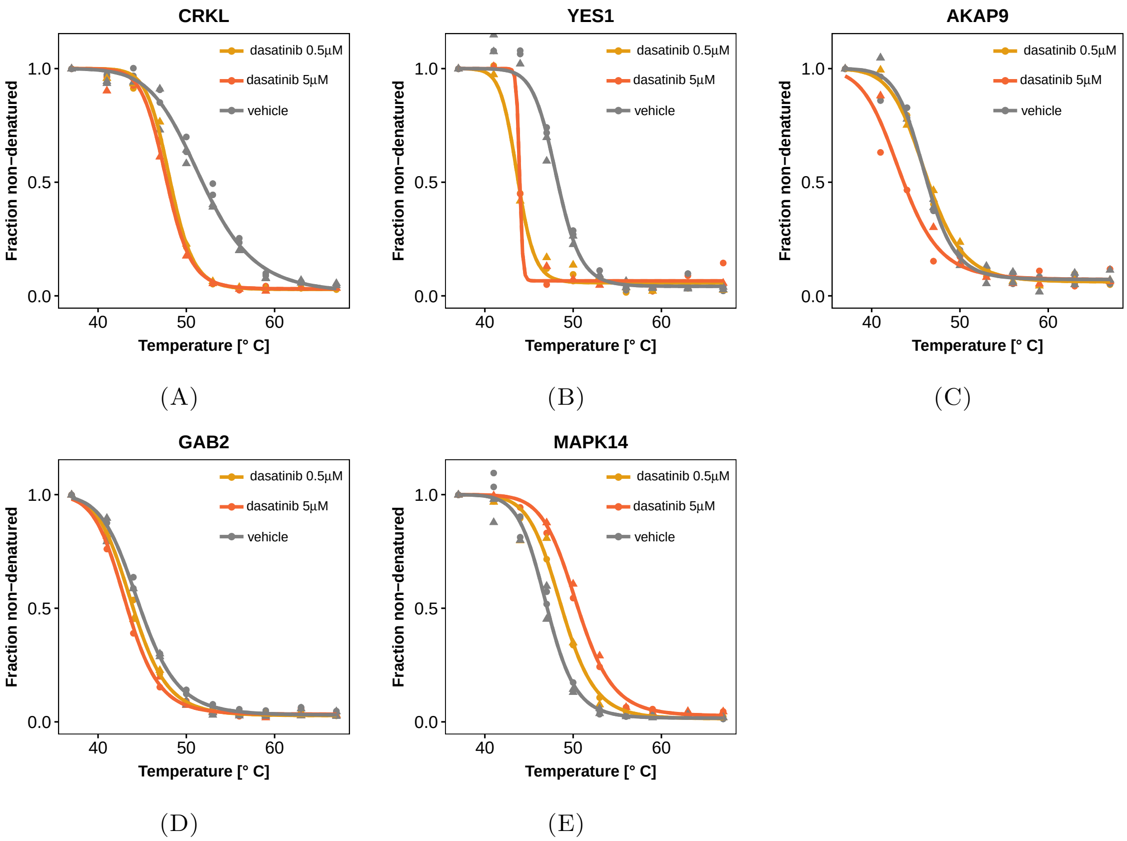


FIGURE S4. The NPARC approach provides the flexibility to simultaneously compare multiple experimental conditions as illustrated by the treatment with dasatinib at two different concentrations. **(A) - (B)**: Crk-like protein (CRKL), and Src family tyrosine kinase (YES1) are interaction partners of the fusion protein BCR-ABL and are destabilized already at small concentrations of dasatinib. **(C) - (E)**: AKAP9, GAB2 and MAPK14 show subtle, dose-dependent but highly reproducible effects on the thermostability after dasatinib treatment. All proteins were detected with Benjamini-Hochberg adjusted F-test p-values $\leq 0.01$ (Supplementary Table S1).

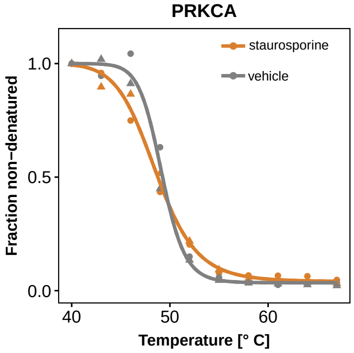


(A)

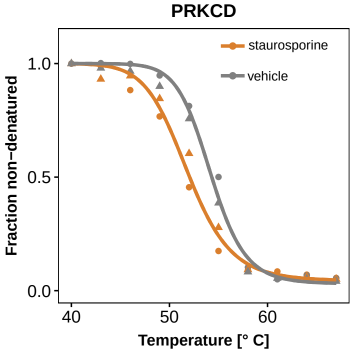


(B)

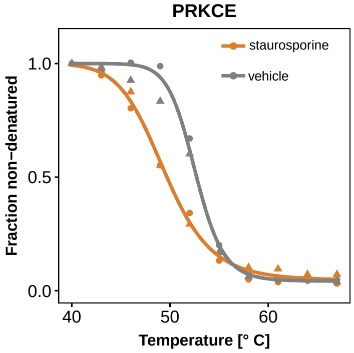


(C)

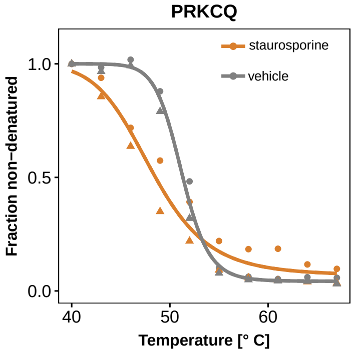


(D)

FIGURE S5. Further examples of the protein kinase C family for which treatment effects are poorly reflected by shifts in $T_{\mathrm{m}}$. Similar to PRKCB (Fig. 1C), the strongest destabilizing effect appears to take place at cooler temperatures than the actual $T_{\mathrm{m}}$ value for PRKCD, PRKCE and PRKCQ. All proteins except PRKCA were detected with Benjamini-Hochberg adjusted F-test p-values $\leq 0.01$ (Supplementary Table S1).
