## Supplementary figures and images for "Non-parametric analysis of thermal proteome profiles reveals novel drug-binding proteins"

### Supplementary Figure S6

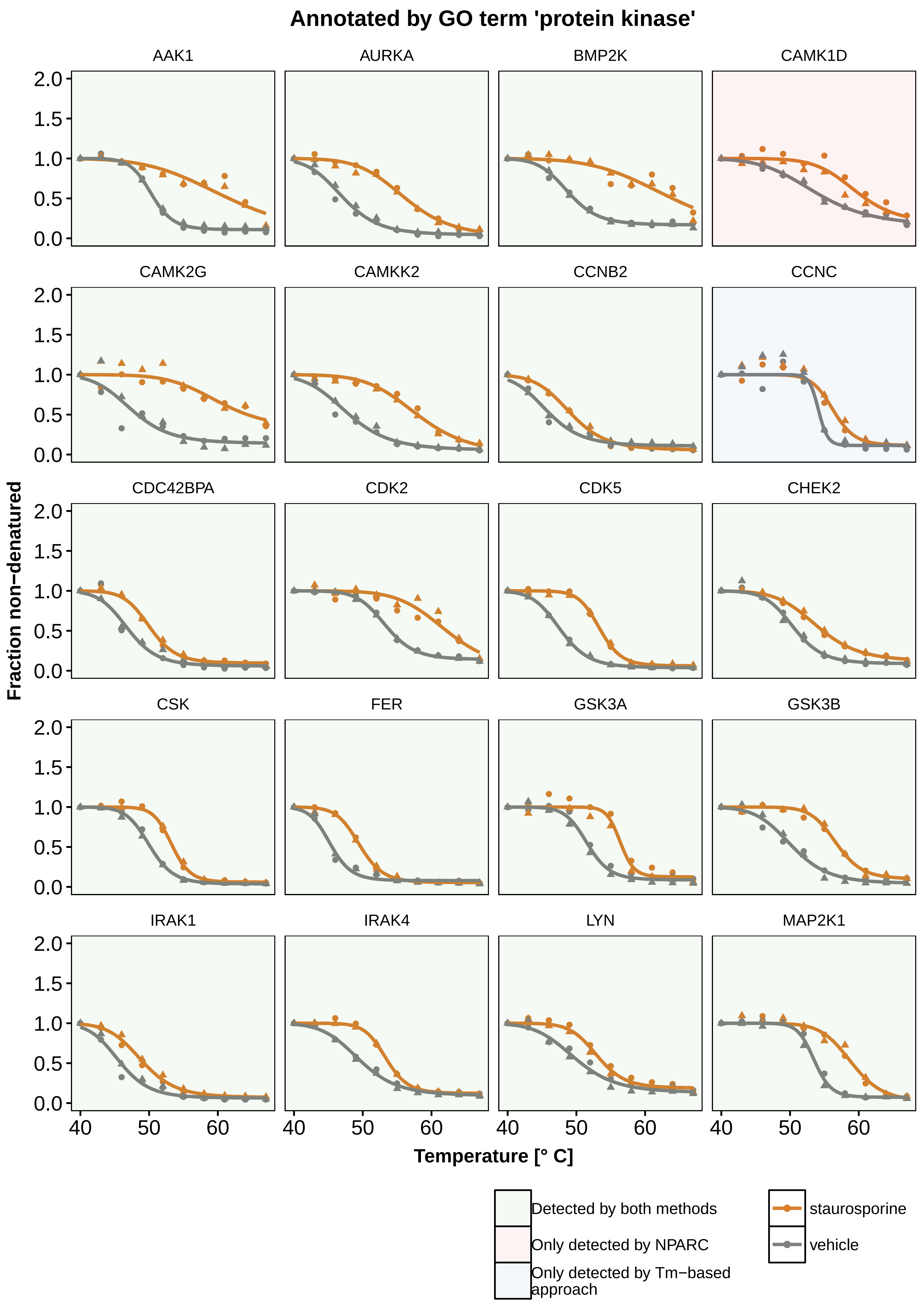

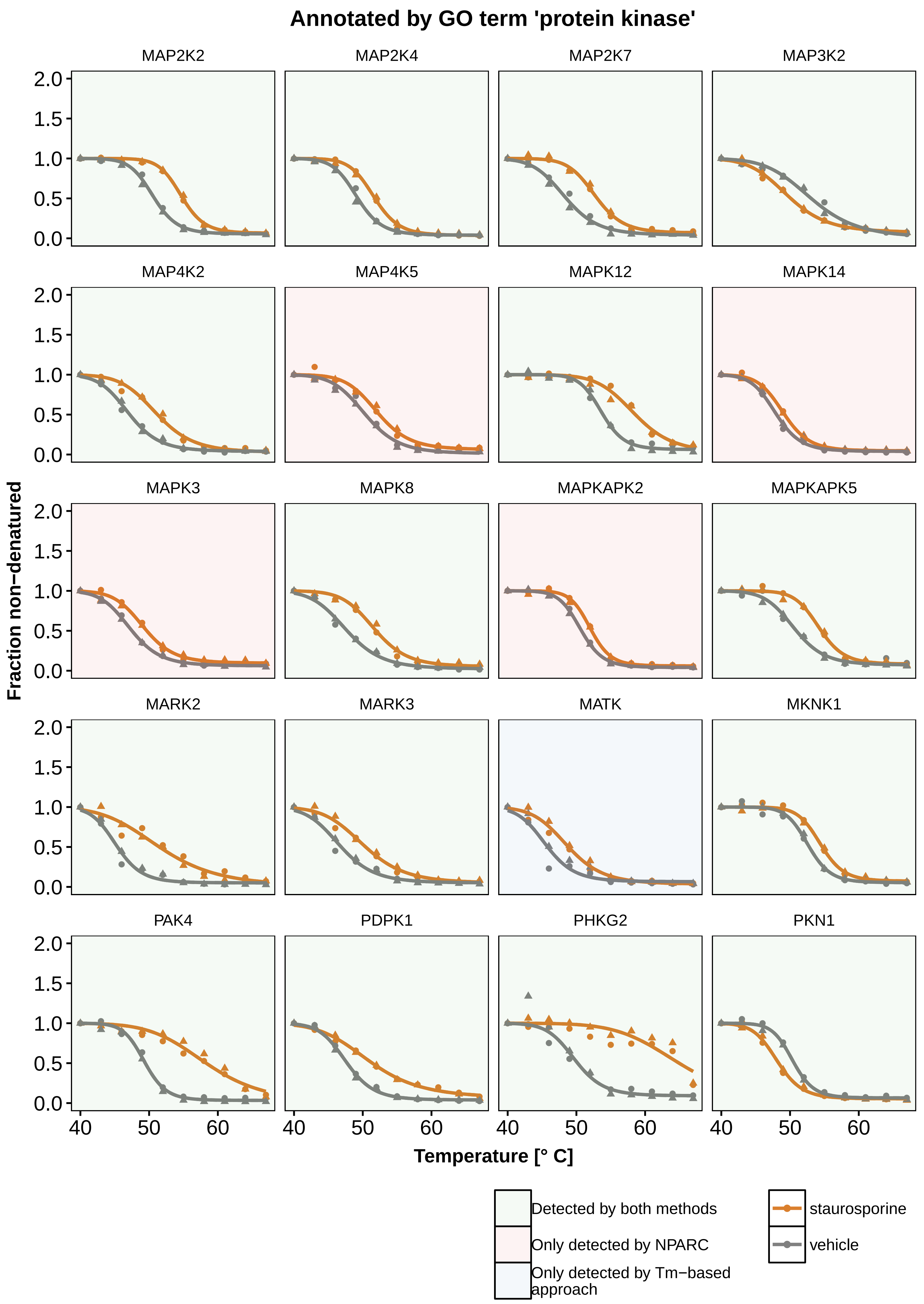

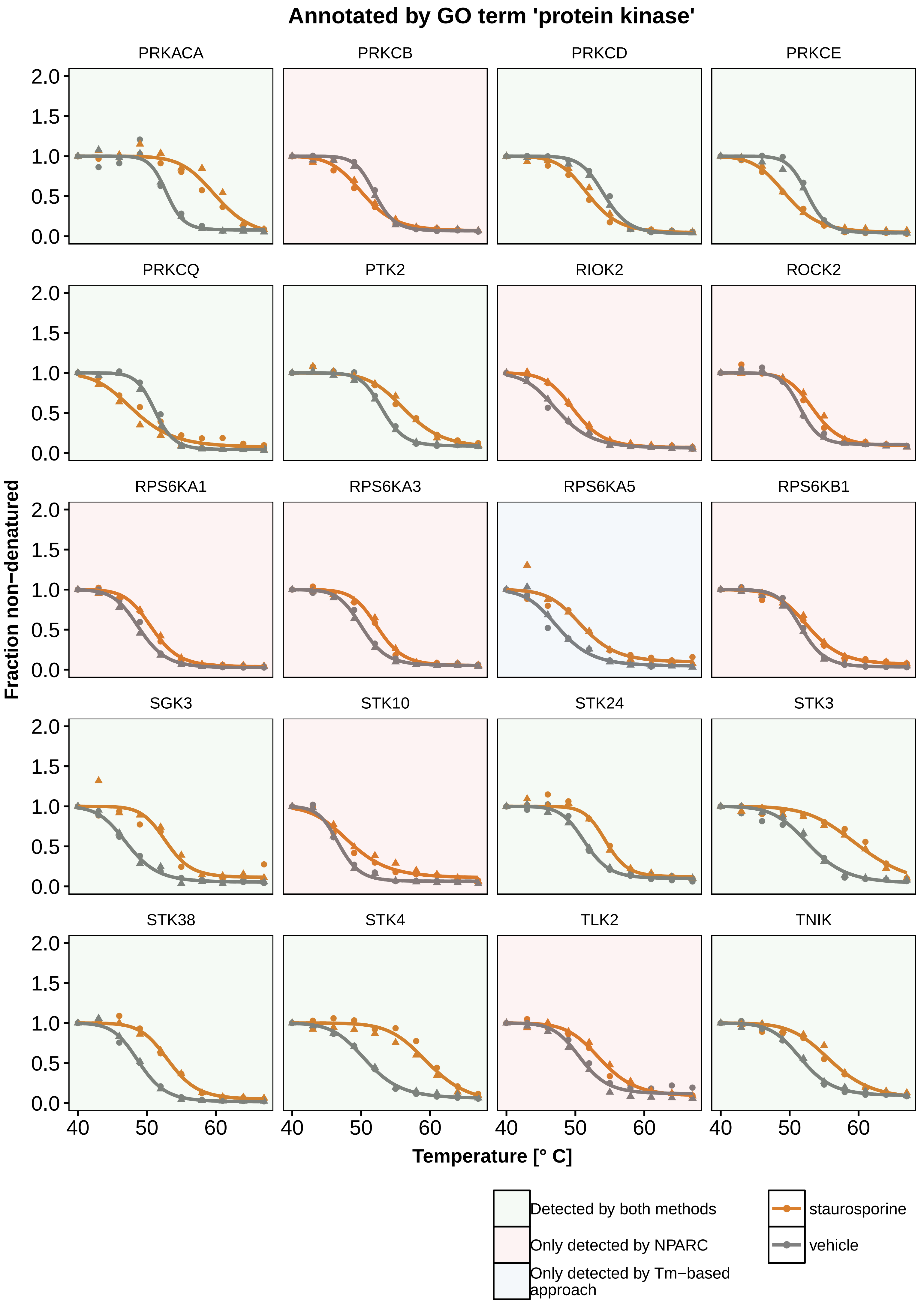

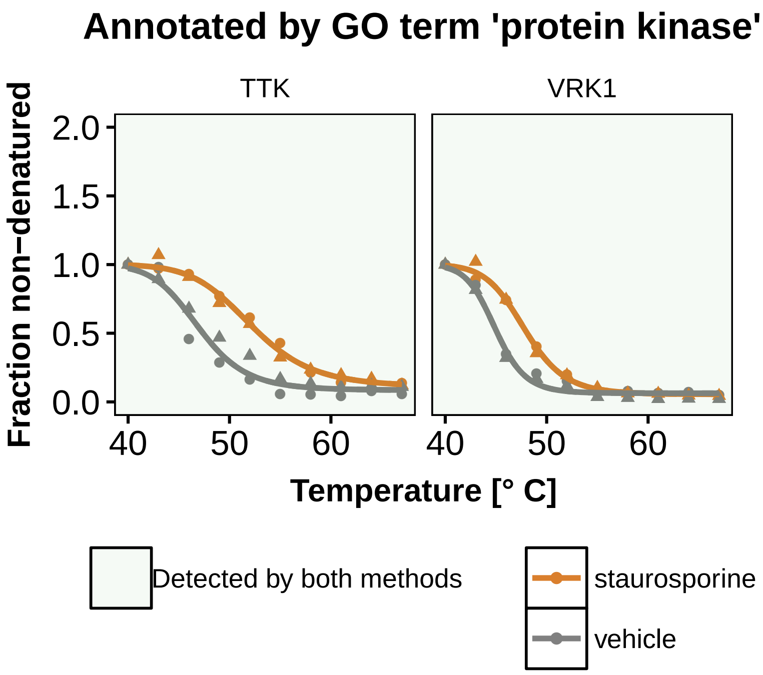

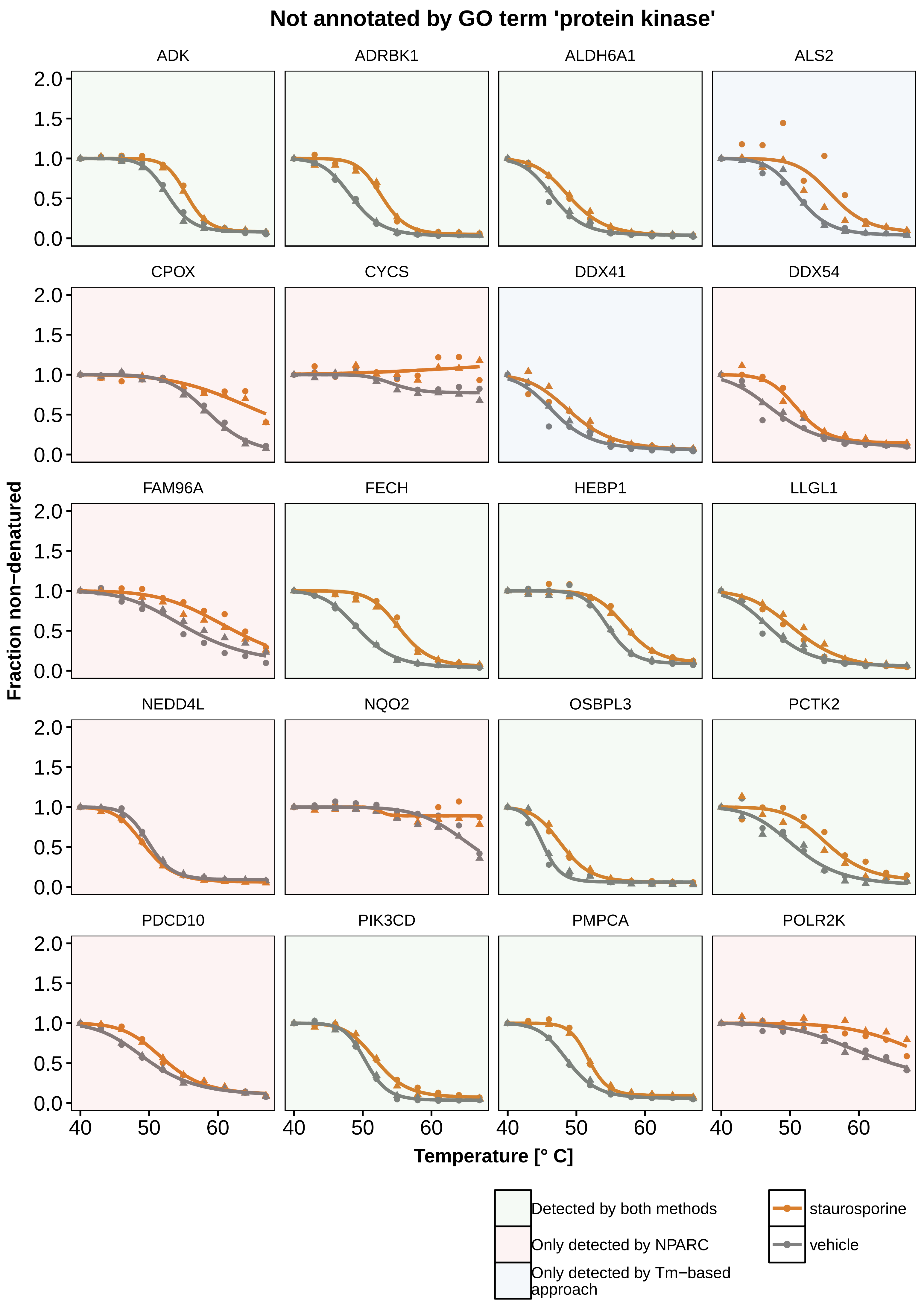

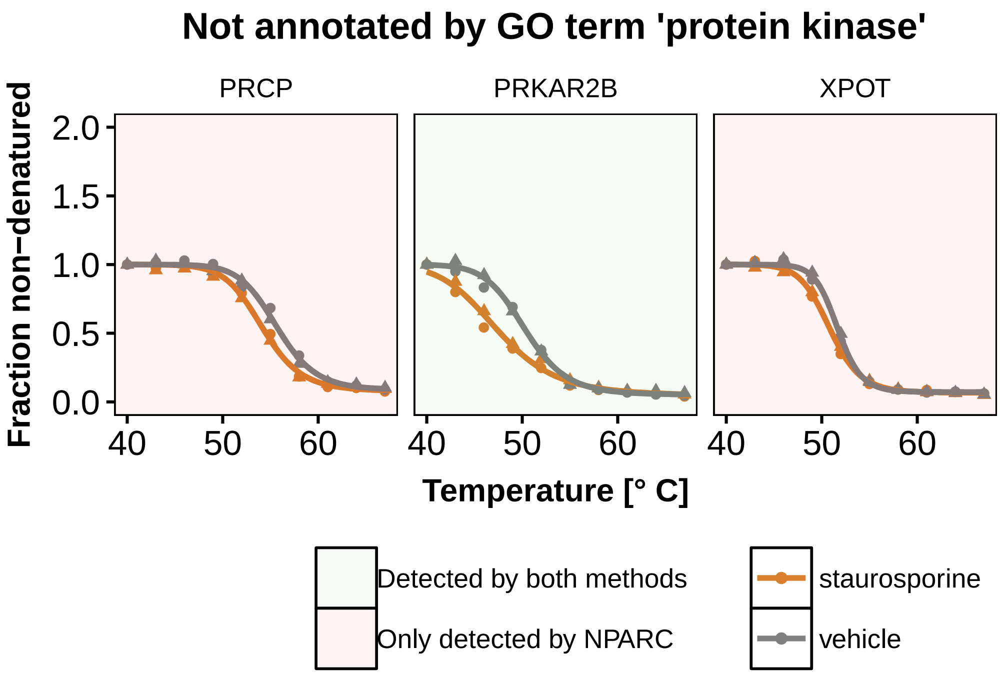

### Supplementary Figure S7

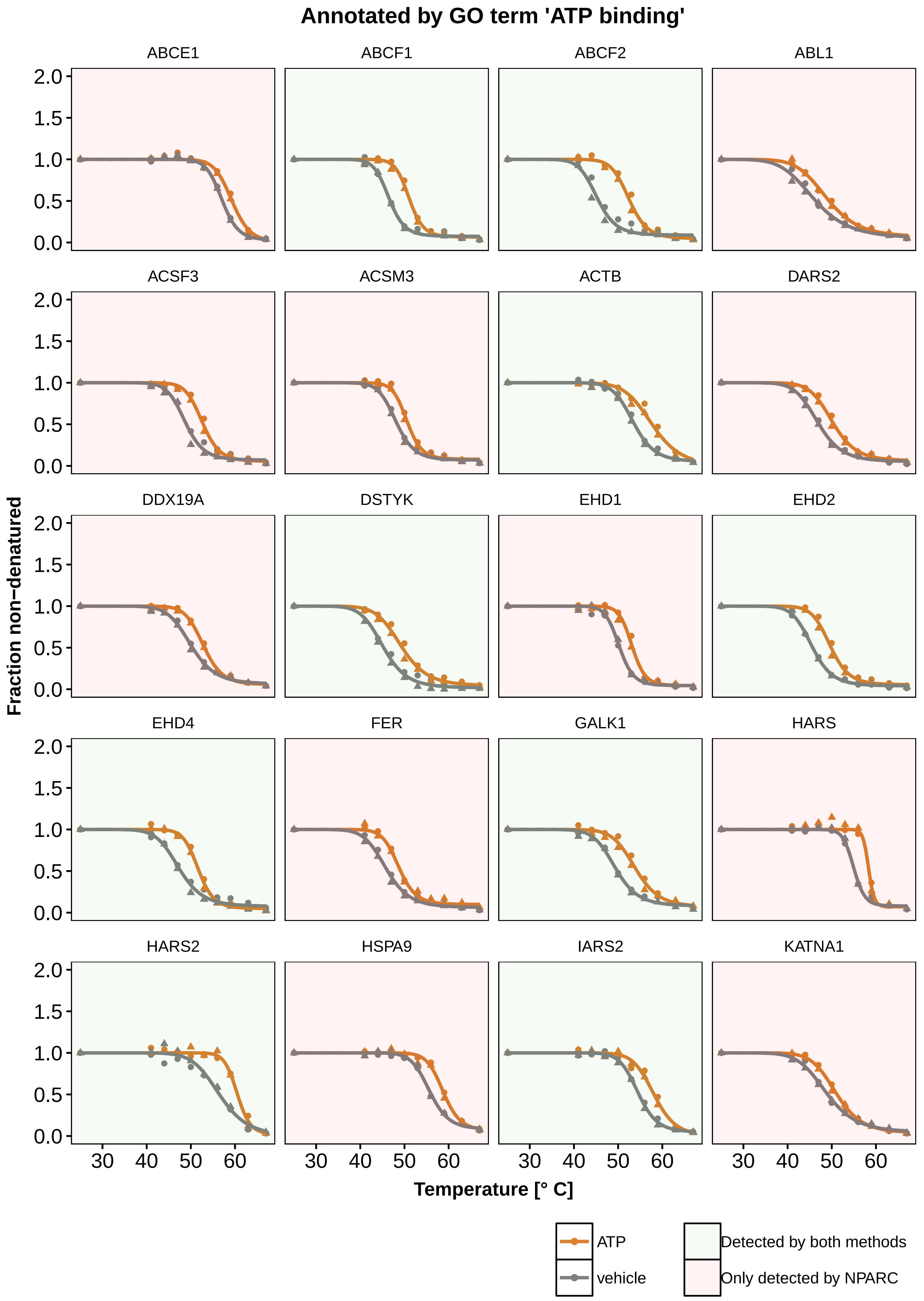

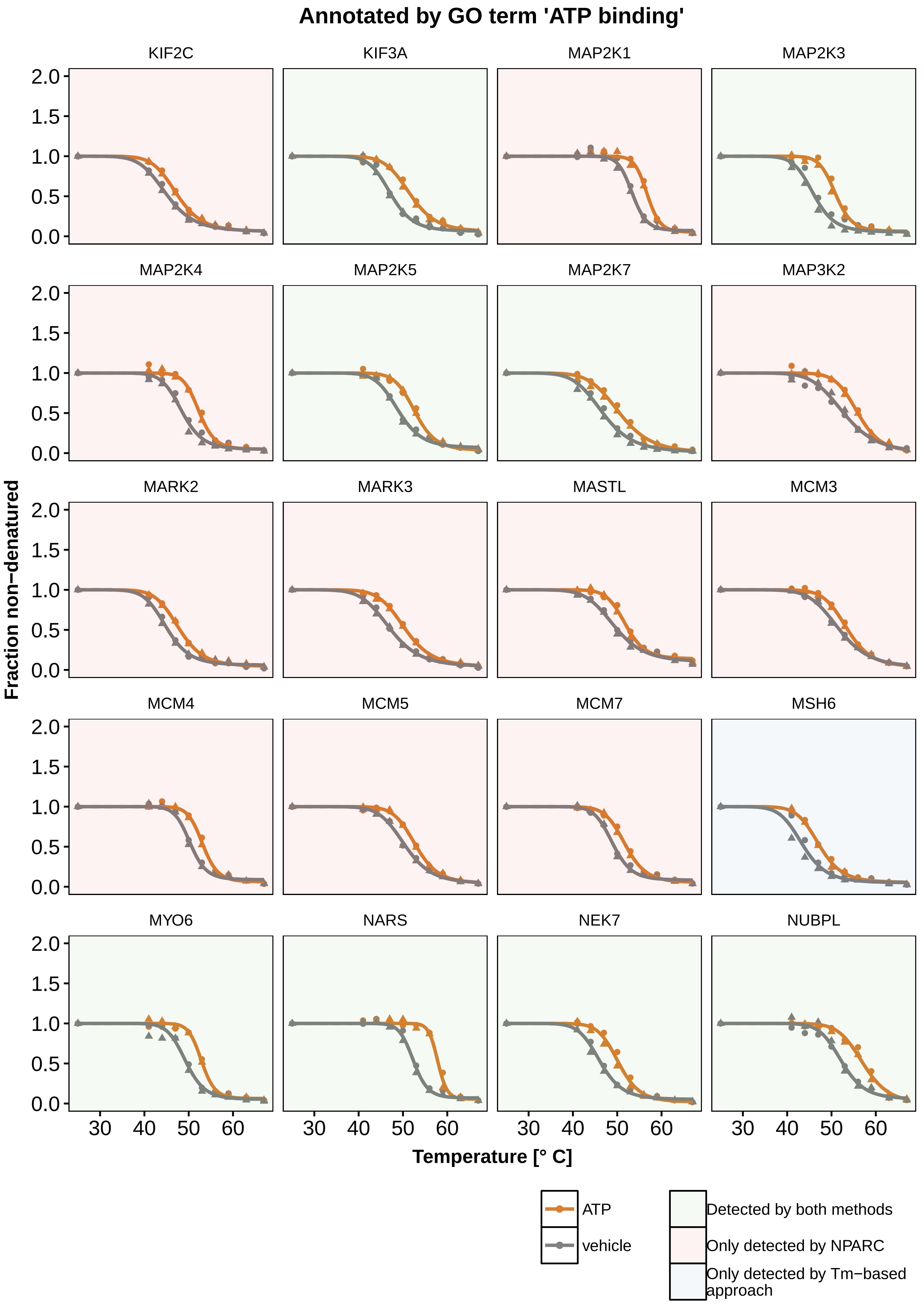

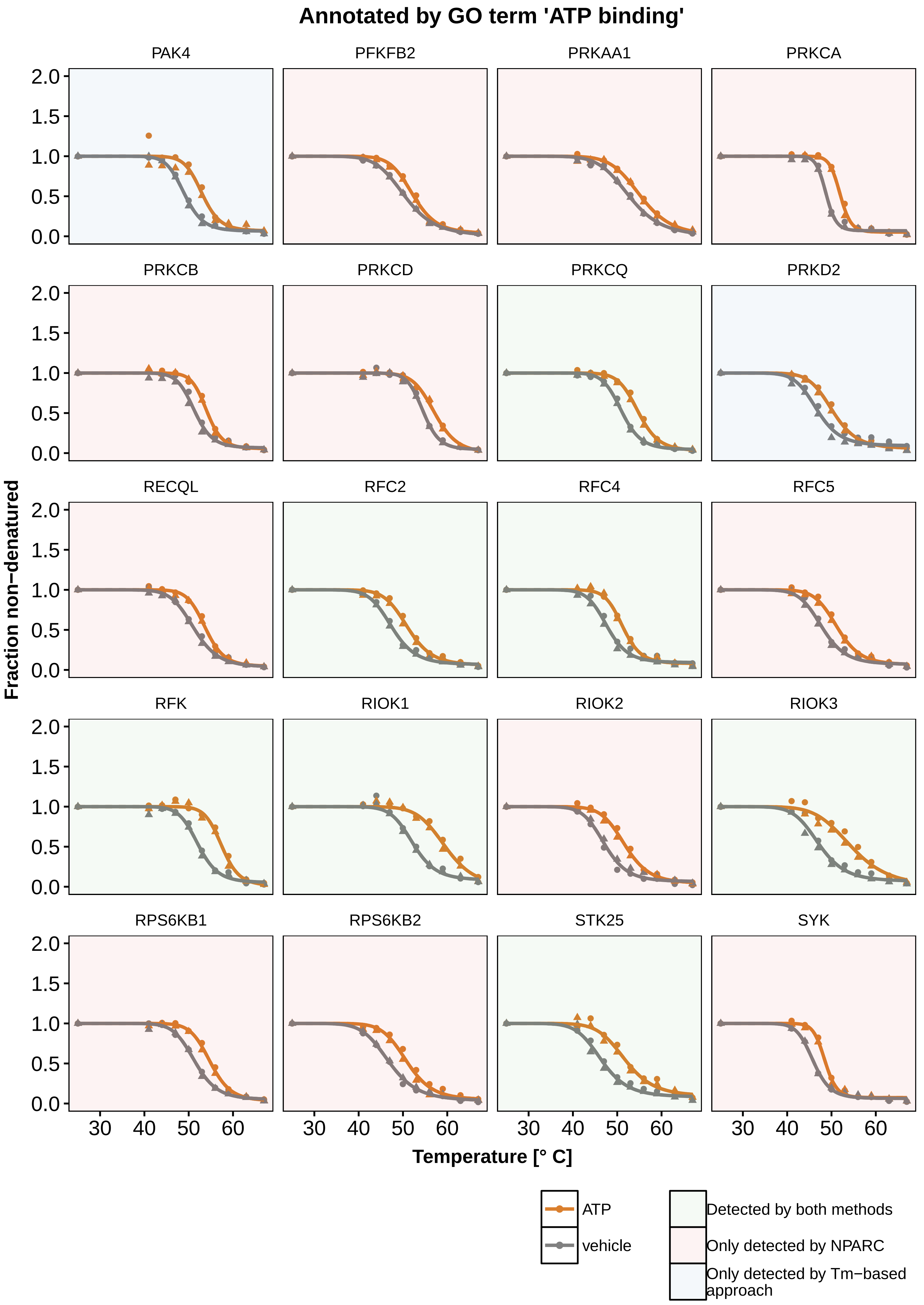

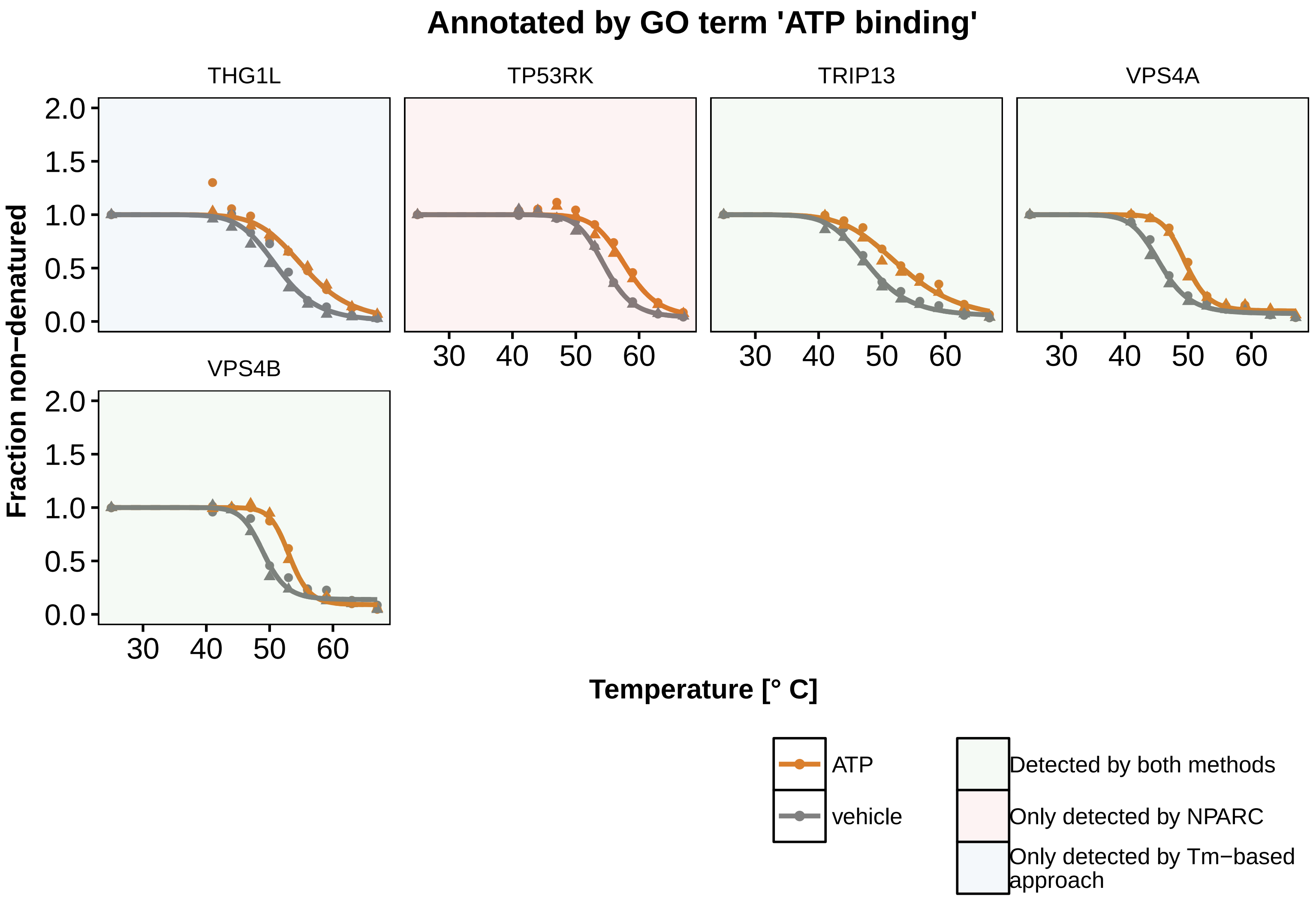

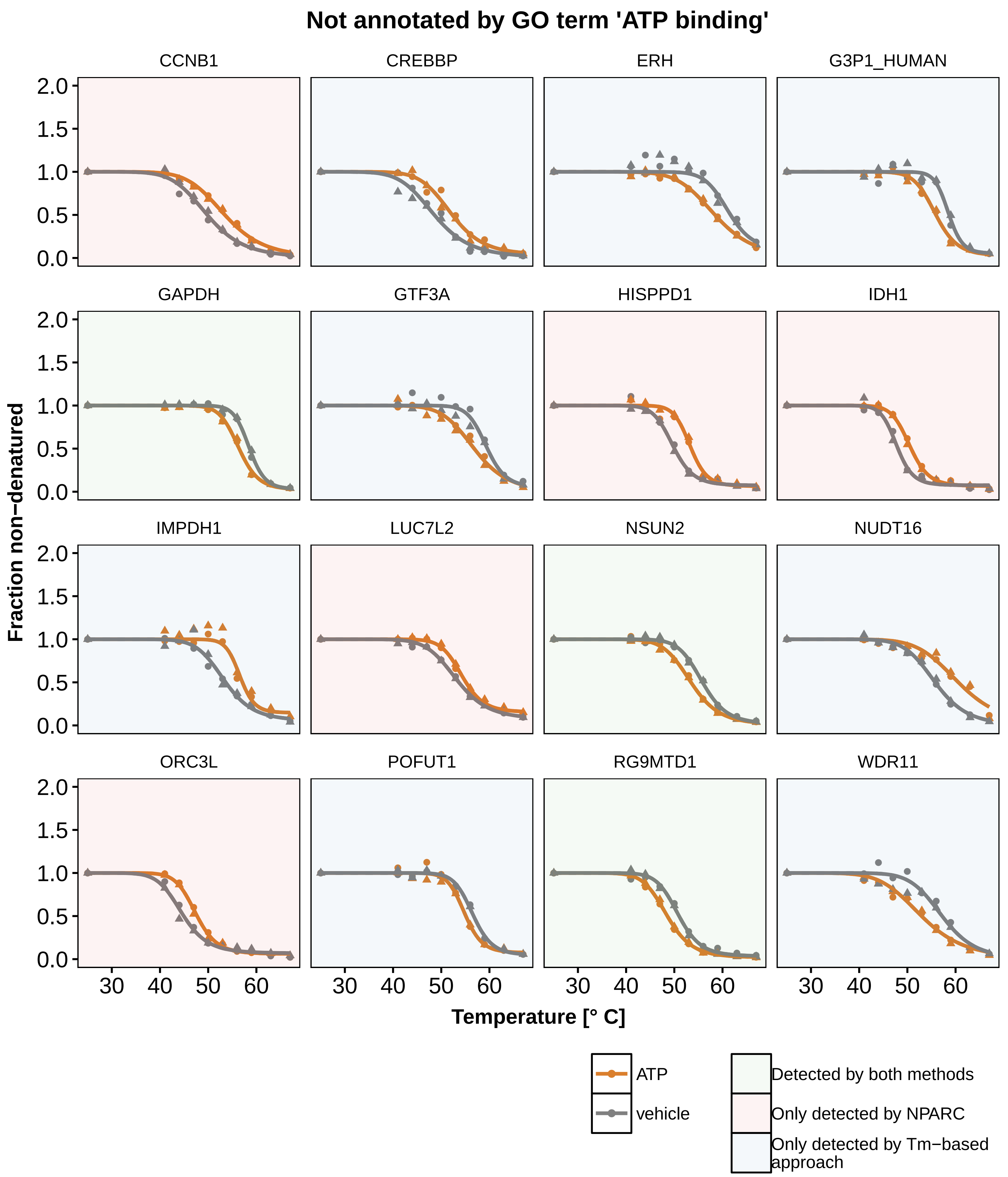
